# Impacts of oil palm plantations expansion on the distribution of terrestrial mammals in South-East Asia

**DOI:** 10.1101/2025.05.14.653968

**Authors:** Michela Pacifici, Andrea Cristiano

## Abstract

The recent expansion of oil palm plantations in Southeast Asia has caused significant range loss for many species. We examined the impact of this expansion on mammal distribution in Indonesia, Malaysia, and Thailand. Using historical range maps (1970s-1980s) and satellite imagery of oil palm plantations (1981 to 2017), we quantified the extent of species’ lost and retained ranges occupied by oil palm plantations before and after 1990. The siamang (+24%), tiger (+19%), and large-spotted civet (+14%) were the most impacted, considering plantations’ expansion in the lost range. All species, including endemic ones, have also seen an increase in their current range. Additionally, we found a significant correlation between the amount of range lost and the area occupied by plantations after 1990. Our findings underscore the urgent need for sustainable land management and conservation strategies to mitigate the adverse effects of agricultural expansion on biodiversity.

## 1. INTRODUCTION

Oil palm (*Elaeis guineensis*) has emerged as the dominant crop in the global vegetable oil industry due to its remarkable land use efficiency and long productive lifespan (Vijay et al. 2016). Native to West Africa, the oil palm tree was introduced to South-East Asia in the early 20th century and has since become a major agricultural commodity. The superior yield of oil per hectare compared to other oilseed crops, such as soybeans or sunflowers, has driven the widespread establishment of oil palm plantations across the tropical forests of South-East Asia (Potts et al. 2014; Dian et al. 2017). The region now accounts for approximately 85% of global palm oil production, with Malaysia and Indonesia as the primary producers (FAO 2019). However, the rapid expansion of oil palm plantations has led to severe environmental consequences, particularly for the rich and diverse ecosystems of South-East Asia. Between 1972 and 2015, Malaysia and Indonesia experienced extensive deforestation driven largely by the conversion of tropical forests into oil palm monocultures (Meijaard et al. 2018). During this period, Malaysia and Indonesia lost respectively 47% and 16% of their forest cover, leading to significant habitat loss and ecosystem degradation and subsequent biodiversity declines (Carlson et al. 2013).

South-East Asia is renowned for its exceptional biodiversity, with a high concentration of unique and endangered species (Hughes 2017). The tropical forests of this region provide critical habitats for a wide range of terrestrial mammals, including iconic species such as the Sumatran orangutan (*Pongo abelii*), the Malayan tiger (*Panthera tigris jacksoni*), and the Bornean pygmy elephant (*Elephas maximus borneensis*). The conversion of tropical forests into oil palm plantations has led to severe declines in mammalian biodiversity, with some estimates suggesting declines of up to 85% in certain areas (Meijaard et al. 2018). For example, the Sumatran orangutan has experienced a precipitous decline in population size, with fewer than 14,000 individuals remaining in the wild, primarily due to habitat loss from deforestation and oil palm expansion (Wich et al. 2016).

Indonesia, in particular, has faced an alarming biodiversity crisis, with mammalian species experiencing extinction rates that are twice as high as those in any other country over the past four decades (Ceballos et al. 2017; Allan et al. 2019). The relentless expansion of oil palm plantations in Indonesia, causing the fragmentation and loss of vital habitats for many species, has been identified as a major driver of this crisis (Dirzo et al. 2014; Ceballos et al. 2015). As a result, numerous mammalian species native to the area are now classified as Critically Endangered or Endangered on the International Union for Conservation of Nature (IUCN) Red List, with habitat loss and degradation acting as the most prominent threats for the persistence of these species (IUCN, 2024).

Despite growing awareness of the environmental impacts of oil palm cultivation (Cazzolla Gatti & Velichevskaya 2020), there is a pressing need for detailed, quantitative studies that assess how changes in oil palm plantation areas specifically affect mammalian species. While previous research has highlighted the broad effects of deforestation on biodiversity (Meijaard et al. 2020), the direct associations between oil palm plantations expansion and changes in the geographic distribution of terrestrial mammals remain relatively obscure. To address this gap, we undertook an analysis of the effects of oil palm plantation expansion on mammalian ranges in South-East Asia. Our study focuses on Indonesia, Malaysia, and Thailand, three countries where oil palm plantations have undergone significant expansion over the past 40 years. By comparing historical (1970s-1980s) and current (up to 2024) mammal range maps, we aimed to quantify the extent of range loss occupied by oil palm cultivation, and to assess its impacts on species distributions.

As global demand for palm oil continues to rise, understanding the specific ways in which oil palm expansion affects mammalian biodiversity is crucial for developing effective conservation strategies. By shedding light on the relationship between oil palm plantation expansion and mammalian range changes, our findings aim to support efforts to balance agricultural development with the preservation of biodiversity and the protection of endangered species. The insights gained from this study will be valuable for policymakers, conservationists, and researchers working to promote more sustainable practices in the oil palm industry and to safeguard the rich biodiversity of South-East Asia’s tropical forests.

## 2. METHODS

To assess the potential impact of oil palm expansion, we consulted the datasets produced by Pacifici et al. (2019; 2023), which contain a total of 475 historic (years 1970s-1980s) range maps for terrestrial mammals at a global scale. These maps were reviewed by species experts and were produced following current IUCN mapping standards and protocols, in order to ensure compatibility with current species maps taken from the IUCN Red List (see SI for details on the methodology). Among those species, we retained only those whose range was included, at least in part, in our study area and for which “Annual & perennial non-timber crops” was listed as a threat in the IUCN Red List. The current range maps (IUCN 2024) were overlaid onto the past range maps to determine the portions of the range that have been lost and retained in the study area over the past 40-50 years. We used GRASS GIS version 7.8.6 (GRASS Development Team, 2020).

To quantify oil palm plantation expansion in Southeast Asia, we used the dataset produced by Danylo et al. (2020). The authors utilized Sentinel-1 imagery to create an oil palm plantation map for Indonesia, Malaysia, and Thailand for the year 2017. This dataset comprises nine tiles of 16-bit GeoTIFFs with a 30-meter resolution, each containing a single attribute value indicating the year the oil palm plantation was first detected since 1980. A value of 4 represents 1984, the earliest year of detection, while a value of 37 corresponds to 2017. We combined the nine tiles using the r.patch function in GRASS GIS and then created two rasters: one for plantations established before 1990 and one for those established after 1990. Given that the Pacifici et al. (2020, 2023) databases include maps spanning the 1970s and 1980s, 1990 serves as the best threshold for identifying changes in ranges possibly attributable to oil palm plantation expansion between the past and the present. We did not account for the specific year of plantation establishment but simply considered the presence or absence of plantations before and after 1990. We overlapped the two plantation maps with the areas of range lost and retained by each species to calculate the percentage of range area occupied by oil palms. We also discussed the limitations of the oil palm database and ran two sensitivity analyses to support our results on plantations expansion (see SI).

We then performed a Wilcoxon test in R to compare the distributions of the percentage of oil palm plantations before and after 1990. This test works by ranking the absolute differences between the paired values, while also considering the direction of the change (whether the percentage increased or decreased). It then sums the ranks of the positive and negative differences separately and uses these sums to calculate a test statistic. The null hypothesis for the Wilcoxon signed-rank test posits that there is no difference between the pre-1990 and post-1990 distributions of the percentages of areas occupied by oil palm plantations. A significant p-value from the test indicates that we can reject this null hypothesis, thereby concluding that there is a statistically significant difference between the two periods (Wilcoxon 1945).

Finally, we tested the relationship between the amount of range lost in the study area and the area occupied by oil palm plantations after 1990 to identify possible correlations between land conversion and species local extirpations. We performed a linear regression and a Spearman’s rank correlation test (Zar 2005), which is a non-parametric test used to test the relationship between two variables when the sample size is small.

## 3. RESULTS

Our final sample included 20 terrestrial mammal species potentially impacted by the expansion of oil palm plantations in Southeast Asia over the past 40 years. The species *Lutrogale perspicillata* did not lose range in the study area, while the species *Rucervus eldii* is now possibly extinct in Thailand, the only country in the study area where the species was present. Therefore, for those species, only one of the analyses on range retained (*Lutrogale perspicillata*) or range lost (*Rucervus eldii*) was present. The analysis on oil palm expansion in the lost range shows significant increases on a variety of terrestrial mammal species (Table 1). Notably, for the siamang (*Symphalangus syndactylus)* only 0.2% of its range lost was occupied by plantations established before 1990; this percentage increased to 23.6% when considering plantations established after 1990. Other species also exhibited considerable increases: the tiger (*Panthera tigris*) from 0.24% pre-1990 to 18.63% post-1990, the large-spotted civet (*Viverra megaspila*) from 0.12% pre-1990 to 14.34% post-1990, the oriental small-clawed otter (*Aonyx cinereus*) from 0.005% to 14.51%, and the dhole (*Cuon alpinus*) from 0.18% to 16.02% (Fig. 1).

**Table 1.**
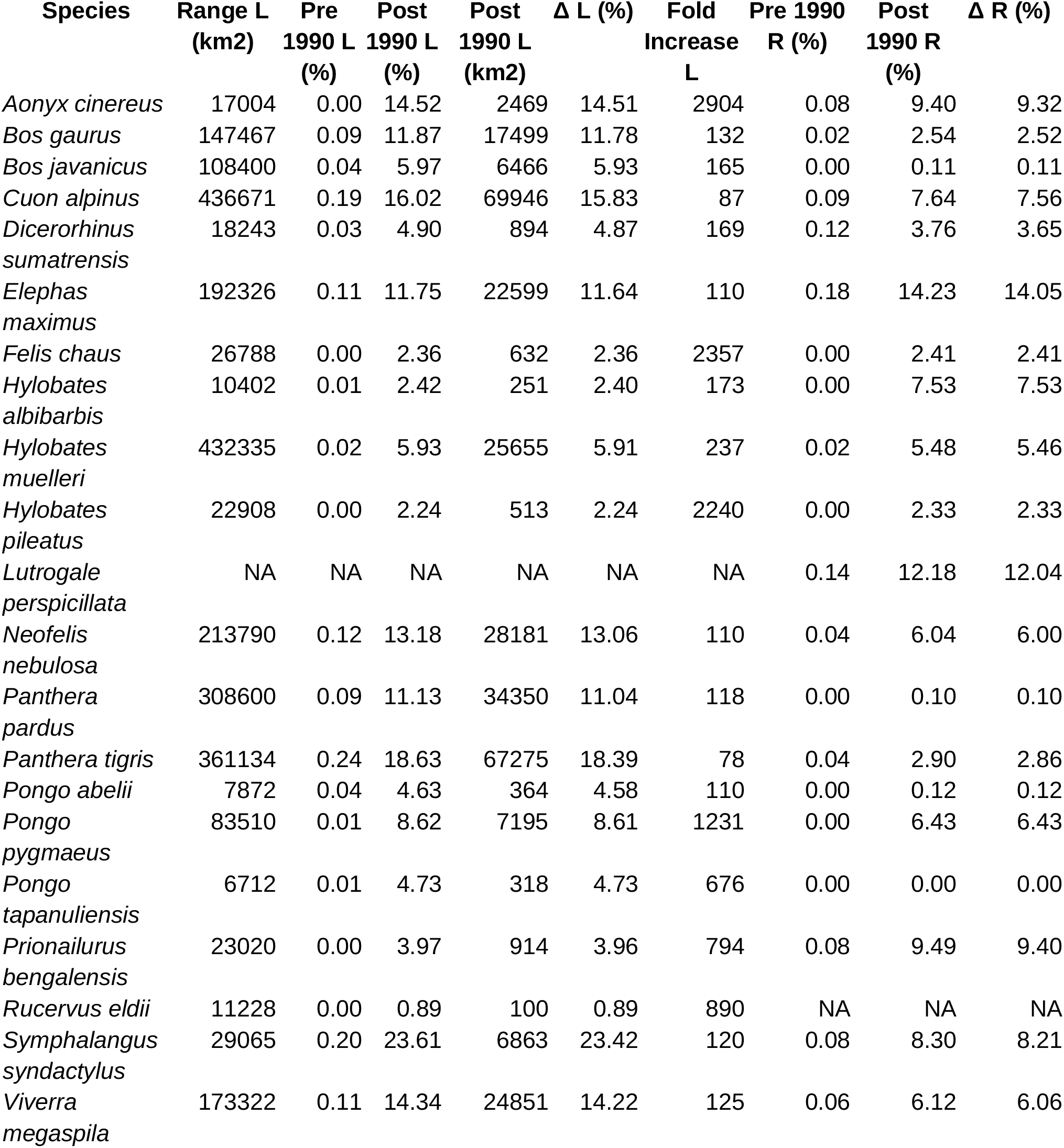
Summary of the percentage of range lost and retained (current) occupied by oil palm plantations before and after 1990. L refers to the lost range and R to the retained (current) range in the study area. Pre/Post 1990 L and Pre/Post 1990 R refer to the percentage of oil palm plantations in the lost and retained areas of the species’ range. Δ is the difference between pre and post 1990 percentages, while Fold increase represents the factor by which the percentage has increased from before to after 1990. All the values have been calculated for the part of the lost and retained range included in the study area. NAs are reported for those species who did not lose or maintain range in the study area.

**Figure 1.**
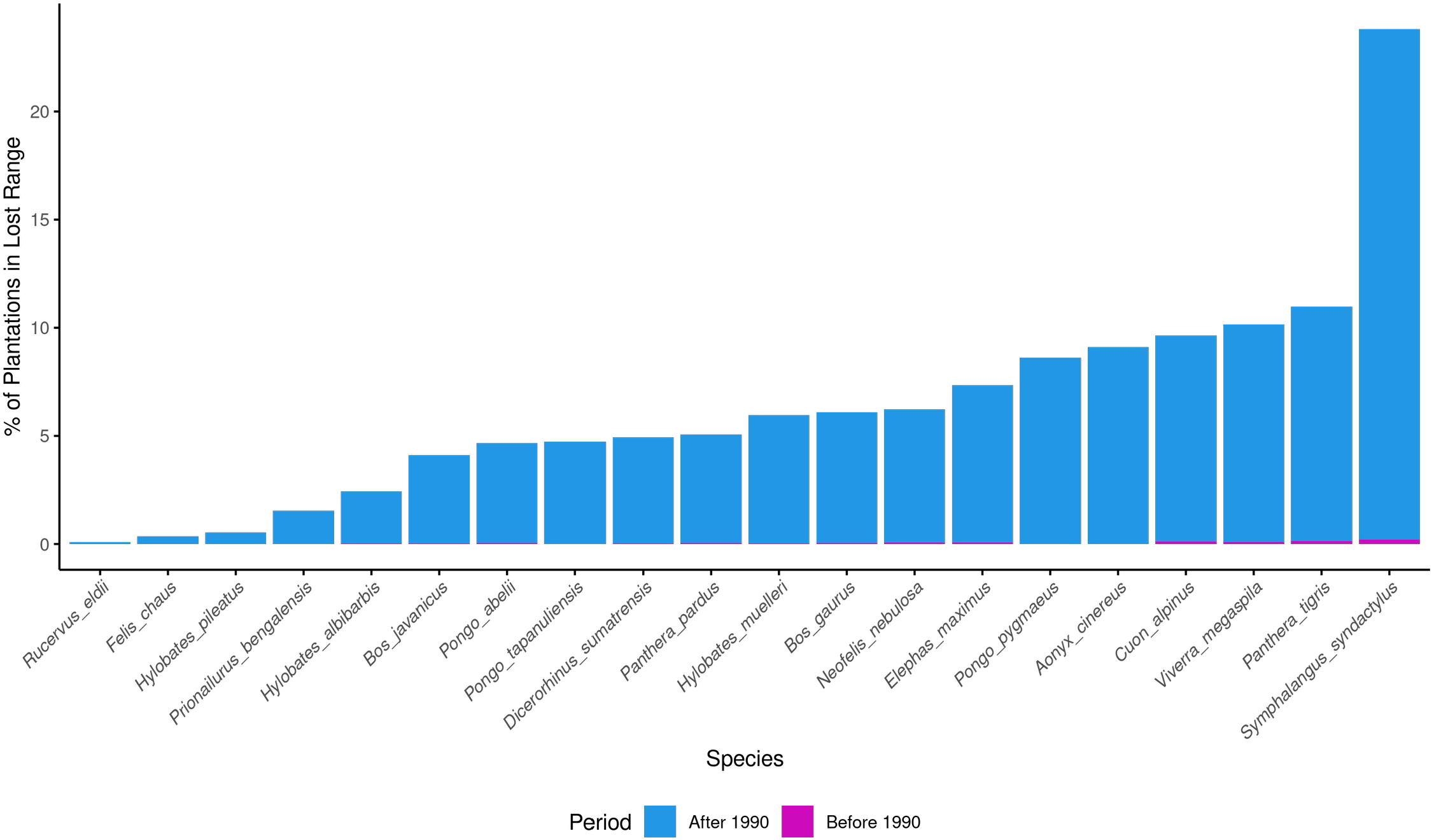
Recent expansion of oil palm plantations in mammals lost range. Stacked bar-chart showing the percentage of the lost range occupied by oil palm plantations before (fuchsia) and after (blue) 1990 in the study area.

All species experienced an increase in the percentage of lost range area occupied by oil palm plantations since 1990 (Table 1). The Wilcoxon test (p-value <0.001) results support the conclusion that there is a significant difference in the increasing coverage of oil palm plantations within portions of species’ range that were lost between 1990 and today. We also found that, on average, 41% of cells with a new plantations have been deforested in the previous 5 years (Table S2).

When analysing the percentage of oil palm plantations within the current range found in the study area, half of the species show values between 6% and 14%, including the Bornean orangutan (*Pongo pygmaeus*; 6.4%), the Bornean white-bearded gibbon (Hylobates albibarbis; 7.5%), the dhole (7.7%), the siamang (8.3%), the Asian small-clawed otter (*Aonyx cinereus*; 9.4%), the mainland leopard cat (*Prionailurus bengalensis*; 9.5%), the smooth-coated otter (*Lutrogale perspicillata*; 12.2%) and the Asian elephant (*Elephas maximus*; 14.3%). For the species endemic to the region considered in this work - *Hylobates muelleri* (5.5%), *Hylobates albibarbis, Pongo pygmaeus*, and *Symphalangus syndactylus* - these percentages reflect the proportion of their entire range currently occupied by oil palm plantations (Table 1).

Finally, we found a significant positive correlation between the extent of range area lost by mammal species and the extent of land converted to oil palm plantations after 1990 (the p-value of the linear model and Spearman’s rank test were both <0.001).

## 4. DISCUSSION

### 4.1 Dramatic increase in oil palm plantation coverage since 1990

Our analysis underscores a dramatic increase in the proportion of oil palm plantations within the lost ranges of the study species from before to after 1990. This significant increase illustrates a profound shift in landscape configuration, as a good portion of oil palm plantations have increasingly encroached into previously unimpacted areas of the species’ ranges (Table S2). The expansion of oil palm plantations reflects a broader trend of intensified agricultural development across Southeast Asia, which has been linked to severe environmental impacts including habitat loss, biodiversity decline, and ecosystem degradation (Meijaard et al. 2018).

For the siamang (*Symphalangus syndactylus*), the increase in plantation coverage from 0.2% pre-1990 to an alarming 23.6% post-1990 signifies a potential extreme transformation of their habitat. This dramatic shift highlights the accelerated pace of agricultural expansion in the region. Such a high percentage of land conversion can have dire consequences for the siamang, whose populations are already facing threats from habitat destruction and poaching (Nijman et al. 2020). The encroachment of oil palm plantations into their range not only reduces the available area but also increases habitat fragmentation, which can isolate populations and disrupt critical ecological processes necessary for their survival, such as foraging and breeding (Miyamoto et al. 2014).

Similarly, the tiger (*Panthera tigris*) and the large-spotted civet (*Viverra megaspila*) have experienced significant increases in oil palm plantation coverage within their lost ranges. For the tiger, the coverage of oil palm plantations within its historic range increased from 0.2% to more than 18.6%. As tigers require extensive and contiguous forest habitats for their large territory and prey base, this increase in plantation coverage represents a possible substantial alteration of habitat conditions, as it reduces prey availability and subsequently increases human-wildlife conflicts, posing a direct threat for the long-term persistence of the species (Dinerstein et al. 2007).

Although the oriental small-clawed otter (*Aonyx cinereus*) and the dhole (*Cuon alpinus*) might exhibit some level of adaptability to habitat changes compared to larger carnivores, the loss of suitable habitats and prey resources due to the expansion of oil palm plantations still poses significant risks. For instance, the reduction in riparian forest habitats can lead to declines in aquatic prey biomass for the otter, while the dhole faces challenges related to both habitat loss and increased competition for prey (Kamler et al. 2015; Wright et al. 2021).

While our study does not directly measure the specific impacts of oil palm plantation expansion on habitat quality and quantity, the observed trends suggest a pattern of deforestation and range fragmentation that is likely to have had significant ecological consequences, possibly contributing to local extirpation in the region. Future research should aim to quantify these impacts by examining changes in species composition, the loss of functional and structural habitat connectivity, and the relationships between habitat loss and population dynamics.

### 4.2 Extent of oil palm plantations within current ranges

Several species currently have significant portions of their distribution covered by oil palm plantations, ranging from 5.5% for the Bornean Gibbon to 14.2% for the Asian elephant. The current proportions of oil palm plantations within species’ ranges indicate that oil palm expansion has encroached significantly into areas that are critical for the persistence of mammal species. For example, the prevalence of oil palm plantations within the Asian elephant’s range may exacerbate habitat fragmentation and escalate human-wildlife conflicts, as elephants are prone to raiding agricultural fields, including oil palm plantations (Calabrese et al. 2017).

For the Bornean orangutan, the situation is particularly dire. Despite the establishment of protected areas, a substantial portion of their habitat remains vulnerable to anthropogenic landscape transformation. Historical deforestation rates in Borneo have been alarmingly high, leading to a projection that more than half of the orangutan’s current range could be lost in the coming decades if current trends continue. Much of the orangutan’s habitat falls within commercial forests and areas earmarked for conversion to agriculture, further exacerbating the threat to this species (Wich et al. 2016).

For species endemic to the region, such as the Bornean gibbon, the Bornean white-bearded gibbon, the Bornean orangutan and the siamang, all forest-dependent species, the notable increase in the percentage of their ranges occupied by oil palm plantations since 1990 reflects possible impacts in the whole species’ distribution (Table S2). Endemic species often have more restricted ranges and are more vulnerable to habitat loss and fragmentation compared to more widespread species (Gonçalves-Souza et al. 2020). The expansion of oil palm plantations into the ranges of these endemic species not only threatens their habitats but also jeopardizes their long-term survival.

### 4.3 Balancing development and biodiversity conservation

Although we recognize potential overestimation of the losses for some species (see SI), our study highlights the urgent need for policy interventions to address the rapid expansion of oil palm plantations in Southeast Asia. The rapid conversion of natural habitats into monoculture plantations has dire consequences for biodiversity, leading to habitat loss, fragmentation, and degradation. The significant increase in plantation coverage since 1990, driven by economic pressures and observed across multiple species, underscores the necessity for effective conservation measures that balance development with environmental protection.

Integrating biodiversity considerations into land-use planning is crucial. Policymakers should enforce stricter regulations on land conversion and promote sustainable agricultural practices. Sustainable palm oil certification schemes have the potential to play a significant role in reducing the environmental impact of palm oil production, but there are still serious concerns about their effectiveness in preventing and mitigating forest loss in tropical areas (Cazzolla Gatti et al. 2019). As oil palm represents a major agricultural commodity worldwide, its controversial role in biodiversity change should be addressed by the production and consumption sides both, for instance fostering practices to reduce the usage and waste of oil palm-based products. Governments, non-governmental organizations and the private sector are called upon to develop innovative solutions that can balance economic growth with ecological sustainability, understanding that trade offs will be unavoidable where the persistence of endangered species is at stake. Strengthening international agreements and funding mechanisms can support large-scale conservation projects, facilitate knowledge sharing, and promote best practices.

Our findings call for immediate action to reform policies and practices related to land use and agricultural development. By prioritizing the conservation of natural habitats and integrating ecological considerations into economic planning, we can ensure the long-term survival of Southeast Asia’s unique wildlife. This approach is essential not only for preserving biodiversity, but also for maintaining ecosystem services that support human well-being. Balancing economic development with environmental preservation represents a fundamental step towards achieving sustainable development goals.

