## Supplementary Information for "Impacts of oil palm plantations expansion on the distribution of terrestrial mammals in South-East Asia"

**Limitations, data uncertainty and sensitivity analyses**

Despite the overall reliability and quality of the datasets used in our analyses, we recognize the importance of discussing the limitations associated with our basemaps. To determine the amount of range lost by a species, we utilized the databases by Pacifici et al. (2019) and Pacifici et al. (2023), which contain range maps for the 1970s and 1980s for 475 terrestrial mammal species. These databases cover just under 10% of terrestrial mammal species and include most of the mammalian orders. However, given that these maps have been published only if considered reliable by experts, it is possible that better-known species may be more represented in the sample. Additionally, past range data are often limited and less comparable to current range maps. To address this, Pacifici and colleagues consulted experts from the current IUCN Red List, who also created the species' range maps, to evaluate the accuracy and reliability of historical maps or to create new maps themselves. They followed the IUCN Mapping Standards (IUCN, 2018) to ensure consistency between past and current maps.

One caveat of the dataset we used to quantify the expansion of oil palm plantations in Southeast Asia (Danylo et al., 2020) is that, particularly in some areas, the presence of false positives may lead to an overestimation of the actual extent of oil palm plantations. To assess the sensitivity of our results to false positives, we used the point dataset employed by Danylo and colleagues to validate their oil palm plantation maps, which contains 10,303 records evaluated by volunteers who determined whether oil palm was present or absent at each sample location. From the attribute table of the dataset, it is possible to identify whether a given point is a true negative, false negative, false positive, or true positive in terms of oil palm presence. For our sensitivity analysis, we only considered the amount of false positives and true positives, and compared them to assess how much of the area identified as oil palm plantation by Danylo et al.’s models could actually not be a plantation (Table S1). We subtracted this percentage from the proportion of species’ lost range occupied by plantations after 1990 (as most of the validation records used by Danylo et al. are recent, and the pre-1990 percentages are very low, we considered those earlier values to be accurate). Next, we compared the percentage of range lost to plantations before and after 1990, both with and without accounting for false positives. The average difference we observed was 0.9% (Table S1), which does not impact our results.

Another issue with the oil palm dataset is that it does not account for prior land use. Unfortunately, high-resolution historical data do not exist, making it impossible to track land-use changes at the scale of the study area and determine whether the area occupied by a new plantation was previously forest, cropland, or another type of land. Despite this uncertainty, it is possible to determine whether the area where a new plantation was established had been deforested in the years immediately preceding its development. We used Global Forest Change data from Hansen (Hansen et al. 2013; Version 1.5), which characterizes forest extent, loss, and gain from 2000 to 2020 at a 30 m spatial resolution and has a high overall accuracy (Galiatsatos et al. 2020), to assess whether a cell where a new plantation was established had been deforested in the previous 5 years. Given the more limited time span covered by the Hansen data compared to our range data, we were only able to assess deforestation from 2005 to 2017 (Table S2). We found that, on average, more than 40% of oil palm plantations were established in areas that had been deforested within the previous five years. Although this timeframe does not cover the full period of our analysis, it confirms that the expansion of oil palm into previously forested areas has been considerable. This suggests that, for forest-dependent species, the impact of oil palm expansion could have been significant. For the remaining 60% of cells, we were unable to directly assess land conversion due to the lack of fine-scale historical data. However, it is important to note that the concept of range differs from that of habitat, which we could not quantify or map over time. Range maps represent the "current known limits of distribution of a species, including all known, inferred, or projected sites of occurrence" (IUCN 2024). This implies that other land cover types, such as croplands that are not suitable for certain species, may have been included in the range loss calculations. This is a limitation of our study. But it is also important to note that not all species in our sample are strictly forest-dependent (only 7 out of 20 species live exclusively in forests, according to the IUCN habitat classification scheme). Thus, the conversion of other land cover types to oil palm plantations could also have had a significant impact on their habitat.

| **Species** | **False positives L (%)** | **Post 1990 L (%) without false positives** | **Δ L (%)** | **Δ L (%) without false positives** | **Difference Δ L (%)** |
| --- | --- | --- | --- | --- | --- |
| *Aonyx cinereus* | 25.71 | 12.65 | 14.51 | 12.65 | -1.86 |
| *Bos gaurus* | 24.00 | 10.44 | 11.78 | 10.35 | -1.33 |
| *Bos javanicus* | 22.06 | 5.31 | 5.93 | 5.27 | -0.62 |
| *Cuon alpinus* | 20.94 | 14.34 | 15.83 | 14.16 | -1.49 |
| *Dicerorhinus sumatrensis* | 30.65 | 4.15 | 4.87 | 4.12 | -0.72 |
| *Elephas maximus* | 22.99 | 10.40 | 11.64 | 10.29 | -1.24 |
| *Felis chaus* | 0.00 | 2.36 | 2.36 | 2.36 | 0.00 |
| *Hylobates albibarbis* | 20.00 | 2.18 | 2.4 | 2.16 | -0.23 |
| *Hylobates muelleri* | 23.53 | 5.24 | 5.91 | 5.21 | -0.67 |
| *Hylobates pileatus* | 0.00 | 2.24 | 2.24 | 2.24 | 0.00 |
| *Lutrogale perspicillata* | NA | NA | NA | NA | NA |
| *Neofelis nebulosa* | 21.39 | 11.77 | 13.06 | 11.65 | -1.29 |
| *Panthera pardus* | 20.62 | 9.98 | 11.04 | 9.89 | -1.05 |
| *Panthera tigris* | 19.15 | 16.85 | 18.39 | 16.60 | -1.54 |
| *Pongo abelii* | 38.89 | 3.73 | 4.58 | 3.68 | -0.86 |
| *Pongo pygmaeus* | 40.00 | 6.89 | 8.61 | 6.89 | -1.72 |
| *Pongo tapanuliensis* | 10.00 | 4.50 | 4.73 | 4.49 | -0.23 |
| *Prionailurus bengalensis* | 0.00 | 3.97 | 3.97 | 3.96 | 0.00 |
| *Rucervus eldii* | 0.00 | 0.89 | 0.89 | 0.89 | 0.00 |
| *Symphalangus syndactylus* | 21.12 | 21.12 | 23.42 | 20.92 | -2.30 |
| *Viverra megaspila* | 14.65 | 13.29 | 14.22 | 13.17 | -0.94 |

**Table S1 Impact of false positive estimates of the presence of oil palm plantation on our models.** L means lost range. “False positives” refers to the percentage of incorrectly identified oil palm plantations in the validation by Danylo et al. (2020). “Post-1990 L without false positives” represents the percentage of range lost to oil palm plantations established after 1990, after accounting for and removing the percentage of false positives. This serves as a conservative estimate. “Δ L” indicates the difference in the percentage of oil palm plantations in the lost range before and after 1990, taking into account potential false presences. “Δ L (%) without false positives” refers to the difference in the percentage of oil palm plantations in the lost range before and after 1990, after removing any potential false presences. “Difference Δ” is the difference between the Δ L values with and without false positives.

| **Year** | **% oil palm plantations in deforested areas** |
| --- | --- |
| 2005 | 97.84 |
| 2006 | 38.59 |
| 2007 | 38.07 |
| 2008 | 39.03 |
| 2009 | 44.44 |
| 2010 | 43.29 |
| 2011 | 44.30 |
| 2012 | 44.55 |
| 2013 | 42.49 |
| 2014 | 40.30 |
| 2015 | 29.90 |
| 2016 | 23.60 |
| 2017 | 12.38 |

**Table S2 New oil palm plantations in areas deforested in the previous five years.** This table shows, for each year of establishment of a new oil palm plantation, the percentage of plantation cells that were deforested in the preceding five years.

Red List Categories and Criteria. Version 16. Prepared by the Standards and Petitions

Committee. Downloadable from https://www.iucnredlist.org/documents/RedListGuidelines.pdf

Pacifici, M., Cristiano, A., Burbidge, A. A., Woinarski, J. C., Di Marco, M., Rondinini, C. (2019). Geographic distribution ranges of terrestrial mammal species in the 1970s. Ecology, 100(7).

Pacifici, M., et al. (2023). Drivers of habitat availability for terrestrial mammals: Unravelling the role of livestock, land conversion and intrinsic traits in the past 50 years. Global Change Biology, 29(24), 6900-6911.
